# Spatio-temporal analysis at landscape scale of the Iberian oak decline epidemic caused by Phytophthora cinnamomi

**DOI:** 10.1101/2020.03.27.011304

**Authors:** Enrique Cardillo, Enrique Abad, Sebastian Meyer

## Abstract

*Phytophthora cinnamomi* Rands is considered a main factor behind the Iberian oak decline (IOD). This epidemic is decimating Holm oaks (*Quercus ilex* L.) and cork oaks (*Quercus suber* L.) which are the keystone trees of a multipurpose, silvo-pastoral and semi-natural ecosystem of 3.1 million hectares in the south west of Europe. Forest diseases are characterized by pronounced spatial patterns, since many of the underlying ecological processes are inherently spatial. To improve the current understanding of such processes, we carried out a complete census of diseased sites via aerial imagery at landscape scale at four different dates over a period of 35 years. We validated our photographic interpretation of *P. cinnamomi* presence in-situ by subsampling soil and roots of diseased sites. To analyse the role of host population heterogeneities in shaping the spread of IOD, we used a ‘self-exciting’ spatio-temporal point process model. Its so-called epidemic component represents the inoculum pressure arising from nearby foci whereas its background component allows for sporadic infections from unobserved sources or disease transmission over larger distances. The best fit was obtained with a lagged power-law for the spatial dispersal kernel, where 49% of the infections triggered by an infected site occur within a distance of 250 meters. Both risk components were found to increase over time. The rate of sporadic infections appeared to be significantly lower in silvo-pastoral systems (dehesas) than in forests and higher in mixed stands and shrub encroached oak-lands. These results may have direct implications for IOD management, for example, the estimated spatial dispersal function helps to define a suitable target area for more efficient control measures. Our results also suggest that silviculture treatments aimed at controlling the density and species composition of oak stands, as well as the abundance of shrubs, could play a key role for disease management

## Introduction

During the 1980s, an epidemic of oak (*Quercus ilex* L. and *Quercus suber* L.) decline and mortality emerged in Spain and Portugal. In the 1990s, the pathogen *Phytophthora cinnamomi* Rands, a primary root pathogen acting on a very wide range of trees and woody plants worldwide, was isolated in the roots of declining oaks in Iberia (Brassier 1996). Since then, *P. cinnamomi* has been very often isolated and its presence reported in declining and dying holm and cork oaks stands (Brasier, 1996; Moreira and Martins, 2005; Rodríguez-Molina, et al. 2005; Serrano et al., 2012; Corcobado et al., 2013). Based on the close association of *P. cinnamomi* with decline sites, its occurrence in diseased roots of Iberian oaks, and its known pathogenicity on holm and cork oak, *P. cinnamomi* has been considered a main factor in the Iberian oak decline (IOD) epidemic (Brassier, 1996; Camilo-Alves et al., 2013).

In Iberia, these evergreen sclerophyllous oaks form a characteristic landscape of open woodlands, known as ‘dehesa’ in Spain and as ‘montado’ in Portugal. Extending over a large area of 3.1 million hectares in Europe, this multipurpose, silvo-pastoral and semi-natural ecosystem is associated with the production of meat, cork, firewood, charcoal, and honey, as well as with hunting and tourism. In ‘dehesas’, trees are considered “ecosystem engineers”, as they allow the maintenance of grass production in poor soils under a semiarid climate (Moreno et. al, 2009). In the ‘sierras’ (low mountain ranges), oaks are also a key species in thicker forests covering the frequently thin soil of mountain slopes jointly with shrubs. In this latter case, human productivity is mainly limited to cork production and hunting. In both ‘dehesas’ and sierras, stands are often jointly populated by the two predominant tree species, *holm* and cork oaks, which form a mixed forest.

Forest diseases are characterized by strong spatial patterns because many of the ecological driving processes are inherently spatial (Ostfeld et al., 2005; Madden, 2006). Beyond the host distribution, spatial patterns may be influenced by the underlying variability in the physical and/or biological conditions favouring the survival and proliferation of the corresponding pathogen, as well as the activity of disease vectors and the presence of reservoirs. While the description and understanding of the epidemic can directly help to develop and to assess the efficiency of control strategies (Madden, 2006), Meentemeyer et al. (2012) point out the paradox that our understanding of disease dynamics occurs at local scales, whereas pathogen invasions and their management occur over landscape scales. Armed with a rich plethora of spatial data, analysis tools and methods, the discipline of forest landscape epidemiology is increasingly focusing on the study of the spatiotemporal dynamics of diseases in complex environmental settings (Holdenrieder et al., 2004; Ostfeld et al., 2005; Meentemeyer et al., 2012).

Relevant concepts to landscape epidemiology include selection of appropriate spatial scale, dynamic modelling, spatially implicit approaches, and selection of ecologically meaningful variables (Meentemeyer et al., 2012). Epidemic analysis is not simply the study of a biological invasion of a given territory characterized by its environmental envelope as well as by human interferences. The most fundamental fact in epidemics is that the habitat of the pathogen, i.e., the host, is itself a living organism (Fodor et al., 2011). Therefore, forest epidemics should be considered invasions that are mainly driven by the interaction of pathogens and host plants populations (Jules et al., 2002; Hodenreider et al., 2004).

Spatio-temporal processes of infectious disease occurrence are characterized by autoregressive or “self-exciting” behaviour (Meyer et al., 2012), where each case or disease event is also a risk factor triggering further cases. This behaviour has been empirically demonstrated in the case of Phytophthora diseases. For example, in the epidemic of root rot (*Phytophthora lateralis*) in Port Orford, cedar populations in Oregon and California, host abundance and proximity to the nearest tree were found to be significant factors associated with an increased infection risk (Jules et al., 2002).

Composition, structure and heterogeneity of forest at stand or landscape scale have been identified as factors that have a significant impact on the abundance of inoculum sources and on pathogen dispersal pathways (Holdenrieder et al., 2004; Plantegenest et al., 2007; Jules et al., 2014). Both are key features in disease epidemiology, but they are poorly characterized for IOD and for other forest diseases. Although host diversity, host connectivity, and host susceptibility have been identified as main factors driving the spread of invasive forest pathogens (Prospero and Cleary, 2017), in communicable diseases a major role is also played by abundance and behaviour of the disease vectors (Fodor, 2011). In turn, these two features depend on stand and landscape characteristics.

Infectious diseases sustained by generalist pathogens, such as *P. cinnamomi*, involve a variety of hosts in a complex, often poorly understood way. It has been generally suggested that a high host diversity is more likely to decrease (rather than increase) the disease risk (Keesing et al., 2006). This reduction is attributed to a dilution effect (susceptible host density reduction) in those habitats where abundant and widespread competent hosts exist and where the pathogen transmission system is frequency dependent (Keesing et al., 2006; Haas et al., 2011; Prospero and Cleary, 2017). In contrast, an amplification effect (risk increase) may occur when the pathogen has a wide range of hosts and when the ecosystem at hand includes abundant reservoir hosts which protect and transmit the pathogen more efficiently than the primary or focal host (Keesing et al., 2006; Prospero and Cleary, 2017). For example, in California forests, *Phytophthora ramorum* causes decline symptoms and mortality in oak and tanoak trees, whereas California bay laurel trees (*Umbellularia californica*) are considered to be relatively tolerant hosts. The bay laurel displays minor symptoms and physiological damages but supports abundant sporulation during the rainy season. In this way, laurel trees play the role of reservoir hosts facilitating pathogen summer survival and reproduction (DiLeo et al., 2009). A similar role is played by *Rhododendron ponticum* as a foliar host for *Phytophthora ramorum* and *Phytophthora kernoviae* in the UK (Denman et al., 2009).

*Phytophthora cinnamomi* is considered a dangerous pathogen, and yet a weak saprophyte that needs living hosts to survive for long periods (Crone et al., 2014). Local abundance of potential reservoirs should increase inoculum pressure and hence the risk of infection of a target plant. In Portugal, after an intensive IOD survey, Moreira et al. (2005) showed that, in addition to oaks, 56% of the surveyed species of shrub flora were infected with *P. cinnamomi*, which was detected mainly in species of the families Ericaceae, Cistaceae and Leguminosae. In Iberian oak ecosystems suffering decline, Costa et al. (2010) have found higher values of mortality in shrublands and open woodlands with shrub encroachment than in ‘dehesas’. In the same vein, Crone et al. (2013) reported that half of the annual and herbaceous perennial plant species living in black gravel infected sites within the *Eucalyptus marginata* (jarrah) forest in Australia, were asymptomatic hosts of *Phytophthora cinnamomi*. In these sites, the removal of vegetation reduced the occurrence of *P. cinnamomi*, thereby enabling their rehabilitation (Crone et al., 2014).

Mathematical modelling of infectious diseases can provide important insights into our understanding of epidemiological processes and has therefore become a key tool for predicting the transmission dynamics in a host population (Maden, 2006; Oli, 2006). Such insights are a key element to guide strategic decisions aimed at controlling the spread of infections (Plantagenest, 2007). In epidemiology, a ‘spatial point pattern’ consists of a set of locations in which a disease event occurred. As soil diseases generally spread by forming discrete foci, it seems reasonable to assimilate the underlying dynamics to that of a spatial point process. However, for communicable diseases, individual events tend to occur in clusters over time, which a simple homogeneous Poisson point process cannot capture. Meyer et al. (2012) developed a spatio-temporal regression framework for the analysis of “self-exciting” point processes. This approach is especially useful to quantify the extent of clustering and to assess the role of environmental factors in disease spread.

The main aims here will be to describe the epidemic of the Iberian oak decline disease at landscape scale and to assess the impact of tree population characteristics on disease spread by estimating their effects on the intensity function of the epidemic process. To fulfil this objective, we proceed in several steps, namely, 1) Detection and mapping of all cases (foci) of oak decline in the study area at four different dates, 2) In-situ confirmation of the relationship between the presence of *Phytophthora cinnamomi* and the more easily observable decline symptoms in oaks, 3) Reconstruction of the space-time dynamics of the disease using an epidemiological model with background and epidemic components and 4) Study of the role of species composition and structure of forest stands in the spread of the disease.

## Methods

### Study area

The study region consists of a rectangular area (29.0 x 18.5 km) located North of the City of Merida, in Spain. Most of its surface (33.895 has) is populated by Mediterranean forests and woodlands of evergreen oaks. The area experiences a Mediterranean climate with a hot and dry summer (monthly average max. temperature: 34º C and precipitation: 7 mm) and a mild winter (monthly average min. temperature: 4º C and precipitation: 80 mm). The soils are nutrient-poor and derived from quartzites and slates in the sierras, slates and granites in the plains and alluvial deposits in the valleys of Lacara and Aljucen rivers. The relief is mainly plain (around 300 m high) with a small sierra in the north (higher peak 570 m) and two river valleys crossing the area in NE to SW direction.

In the plains, these sclerophyllous woodlands have been historically managed as a silvo-pastoral system known as ‘dehesa’. This system integrates pasture, livestock and a sparse population of large oaks. Holm oak is more prevalent than cork-oak, but both species are widely used to supply shadow, shelter, acorns and firewood. In addition, cork oak is debarked every nine years to produce cork stoppers. All year long, the dehesas are grazed by sheep and cows for meat, but during autumn and winter, Iberian pigs are introduced in the ecosystem for being fed with acorns.

In the sierras, trees are accompanied by an understorey of woody shrubs (matorral), mainly composed by *Cistus sp*. In the ‘dehesa’, shrub encroachment is prevented by grazing and periodic disking. Wherever the forest becomes thicker, game is usually abundant, with overstocked populations of deer and wild board at certain sites. Declining holm-oaks and cork-oaks have been observed in the area for more than 30 years, and P. cinnamomi has been repeatedly isolated from both hosts. The pathogen was found for the first time in 1999 in the farm ‘El Rosal de Abajo’ (Rodríguez-Molina et al., 2003), and ten years later P. cinnamomi was detected in four additional sites during a regional survey of oak decline (Corcobado et al., 2013).

### Oak decline detection and mapping

A complete census of incident cases was carried out by photo-interpreting aerial imagery. To detect oak decline, we used high resolution (pixel width: 0.25-0.50 m) aerial digital imagery collected at the study site during spring or early summer at four different dates: 1981, 2002, 2007 and 2016. These mosaicked, radiometrically and geometrically corrected images were supplied by the Cartographic and Territorial Information Centre of Extremadura, (Spanish National Program of Aerial Orthophotography, PNOA 1981-2016 CC-BY 4.0).

Oak decline was detected by a trained human photo-interpreter who assessed the status of oak tree crowns. According to a photo-interpretation pictorial key (Ciesla, 2000), six crown decline stages were considered: healthy, chlorotic, branch dieback, recently dead, older dead trees and snag with only main stem and branches. Wherever mortality and at least three oak crowns exhibiting two or more decline stages were observed simultaneously, an oak decline focus was considered to exist and, consequently, it was mapped. Assistance for diagnosis came from change detection using older imagery as background. A minimum circular polygon was drawn around symptomatic trees and subsequently registered in a GIS database.

Sites disturbed by other well-known stress factors were excluded. For example, stands damaged by wildfires or defoliated by gypsy moth (*Lymantria dispar* L.), stands recently pruned, fenced sites heavily trampled by overcrowded livestock, and trees on reservoir banks subject to strong water level changes were identified, but not mapped.

### Field sampling and Phytophthora cinnamomi isolation

To validate the diagnosis of the photo-interpreter and to confirm the presence of P. cinnamomi as damaging agent, ground truth observations were conducted. A sample of 22 oak stands was selected randomly from a collection of stands where disease foci had been detected upon site inspection during early spring. In each focus, the presence of oak decline symptoms (dead trees, crown transparency and branch dieback, see figure 2) was checked and *Phytophthora cinnamomi* isolation in trees inside foci was attempted by means of laboratory analysis of rhizosphere soil samples (Jung et al., 1996). At each site, three soil sub-samples consisting of monoliths containing oak roots (approximate size 30 x 30 x 30 cm.) of three trees with early symptoms were extracted at a depth of one meter from the stem base. Prior to soil sampling, the organic layer was removed. In the laboratory, soil sub-samples (around 1 litre of soil) from each focus were bulked and mixed thoroughly. Isolations were carried out by flooding soil samples with distilled water and by subsequent baiting with 2-7 days old cork oak floating leaflets. After 3-7 days at 20ºC, discoloured leaflets were cut into pieces and plated onto a selective medium (NARPH, Hüberli et al., 2000). After 24 h at 20-25 ºC in the dark, *Phytophthora* hyphae were transferred onto V-8 agar. *Phytophthora cinnamomi* was identified by its distinctive morphological traits with the help of an optical microscope. In addition, the downslope streams in the vicinity of one of the subsamples (n = 12) of the sampled foci were also baited for *P. cinnamomi* isolation. In early spring, ten mesh bags per site were filled with 2-7 days old Quercus suber leaflets, placed just below the stream water surface, and tied with a nylon rope to the stream bank. After 7-10 days in the water, the baits were retrieved and processed in the laboratory as described above. To confirm morphology identification, the ITS region of ten soil isolates were amplified and sequenced using ITS4 and ITS6 primers. The sequences were subjected to an NCBI BLAST search to find homologies with *P. cinnamomi accessions*.

**Figure 1.**
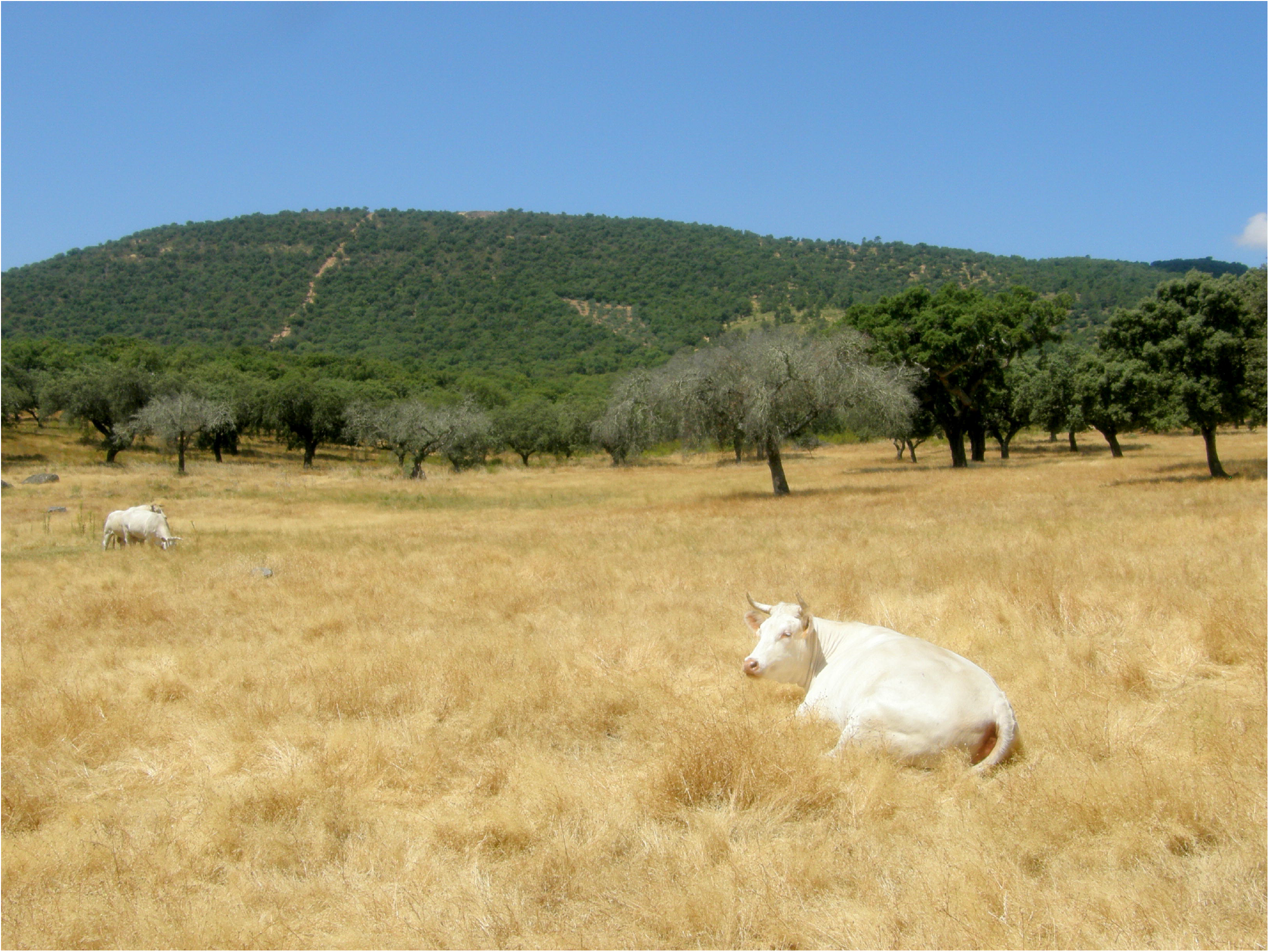
Oak decline focus in a dehesa farm located in the study area. Oak forest and woody crops (olives) can be seen on the background hills.

**Figure 2.**
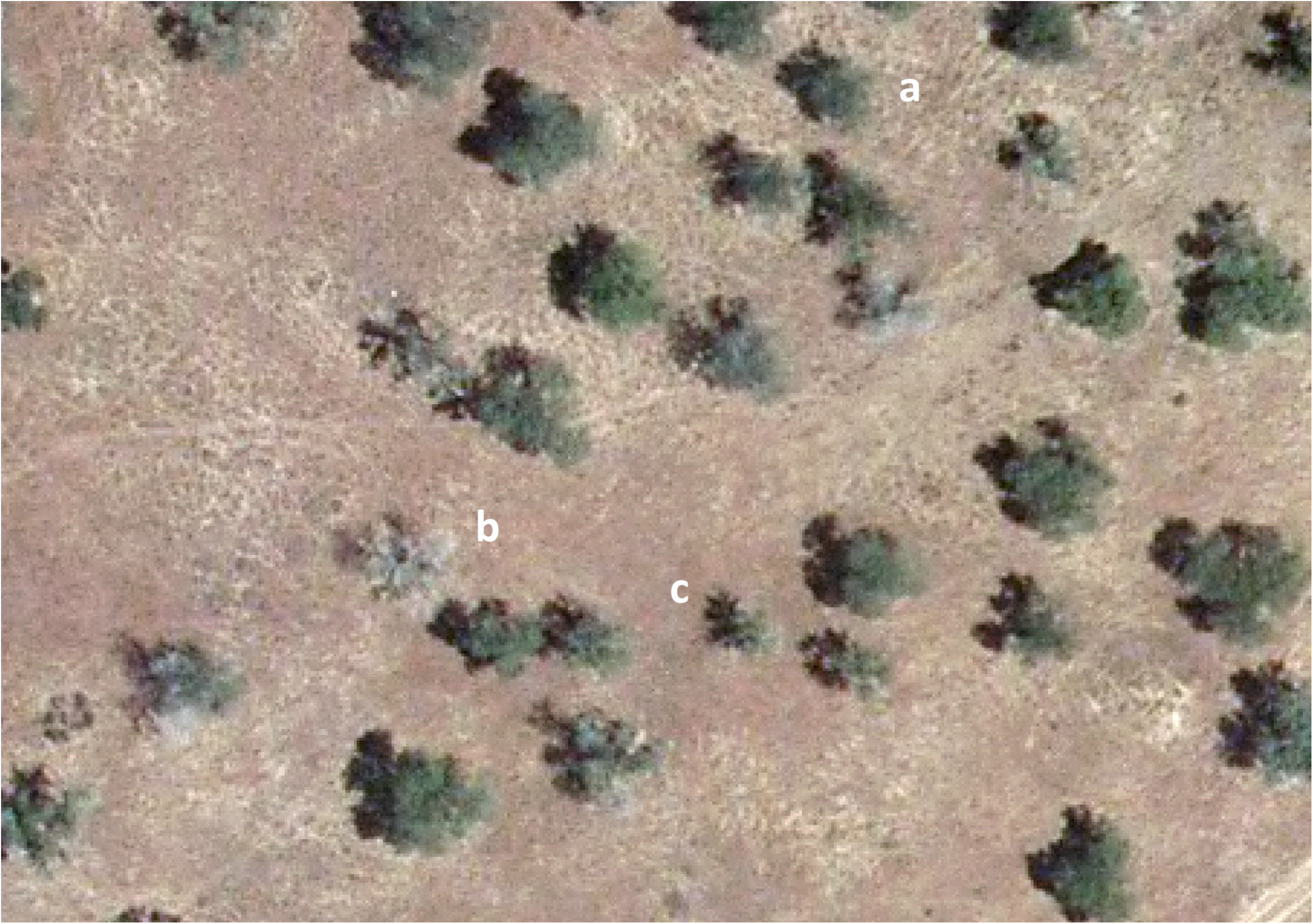
Aerial RGB image of a focus of oak decline in which trees at several stages of decline can be observed: pale green crowns (a) indicate lost leaves, while white crowns (b) reveal sudden death. The crown shadow in the shape of a star (c) is a consequence of branch dieback. (Printed under a CC BY license, with permission from Junta de Extremadura CICTEX, original copyright: Orthoimagery CC-BY 4.0 CICTEX, Junta de Extremadura).

### Oak population covariates

Landscape scale data from tree host populations were obtained from the Forest Map of Spain (FMS; Ministerio de Agricultura Pesca y Alimentacion, 2006). This is a digital database, at 1:50000 scale, consisting of a tessellation of the territory in homogeneous vegetation units (1286 tessellae in this case) with a relatively uniform age distribution, composition and structure. In addition to geographic boundaries, FMS registered relevant information for every forest stand (tessella), including the three main tree species present, their development stages, occupancy (surface proportion occupied by each species), total and tree canopy cover, and forest structure. Using the FMS database, the following covariates were calculated for the 288 oak stands (cf. table 2 and figure 3): tree canopy cover (TCC); shrub canopy cover (SCC) [computed as the difference between total and tree canopy cover]; dominance index (DI), taking values close to zero in stands populated by balanced mixes of oaks species and values close to 5 in pure stands; forest stand use (FSU), a categorical variable which can either be ‘dehesa’ in case of silvo-pastoral use, or ‘forest’ otherwise; and stand connectivity (CONNECT), another categorical variable with either of two values: connected (when stands are conterminous with other forest stands), or isolated, (if stands are surrounded by crops). Dummy effects for the different time periods were also included in the analysis.

**Figure 3.**
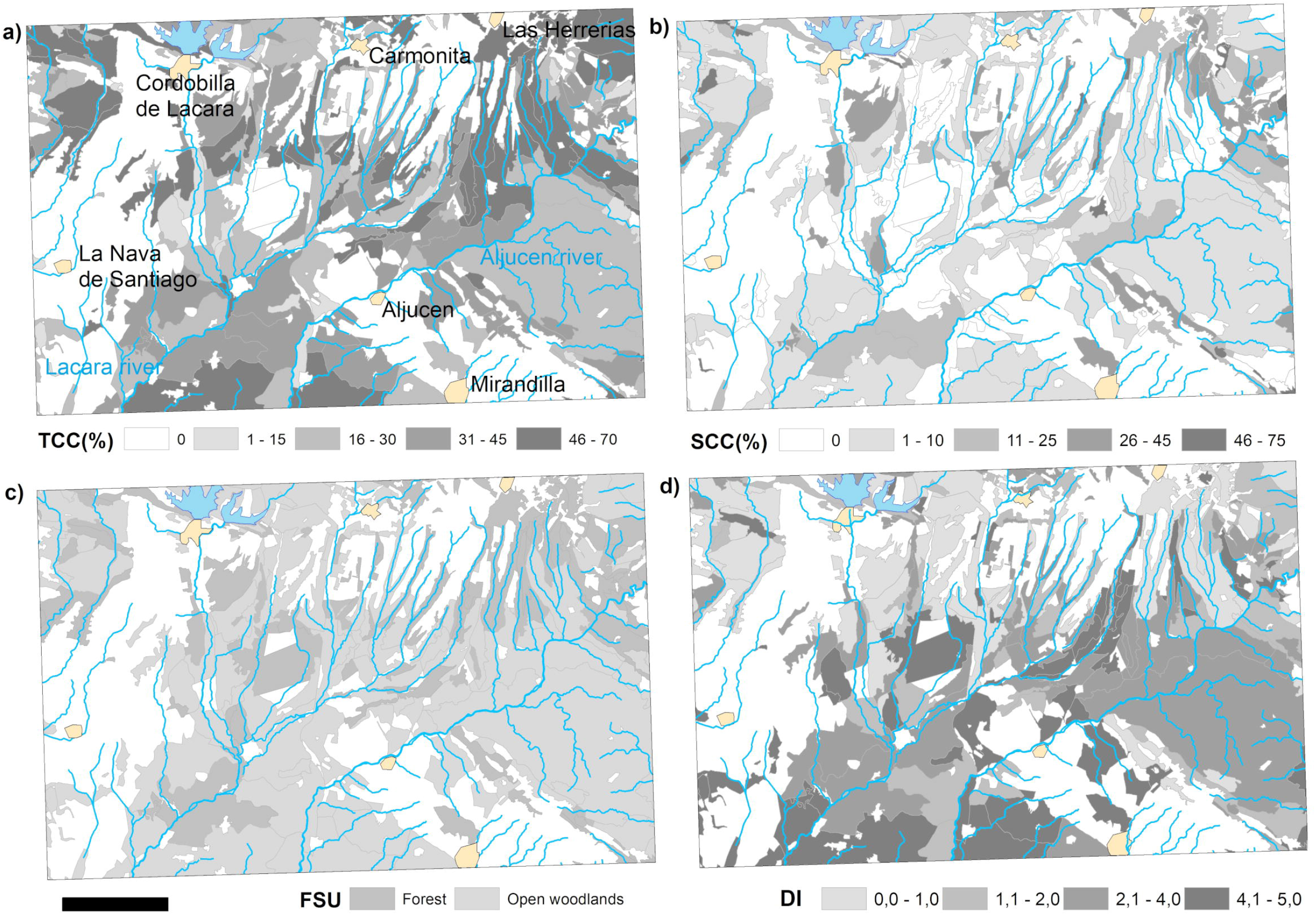
Spatial distribution of covariates obtained from the Forest Map of Spain. (Ministerio de Agricultura, Pesca y Alimentacion, 2006). a) Tree canopy cover (TCC %), defined as the proportion of the forest covered by the vertical projection of plant crowns. b) Shrub canopy cover (SCC %). c) Forest stand use (FSU): Forest or silvo-pasture in open woodlands. d) Dominance index (DI), defined as an indicator of the degree of mixing of tree species in a given stand (the dominance index ranges from a null value for a 50% mix of both species to a maximum value of 5 for a pure stand of either species).

**Figure 4.**
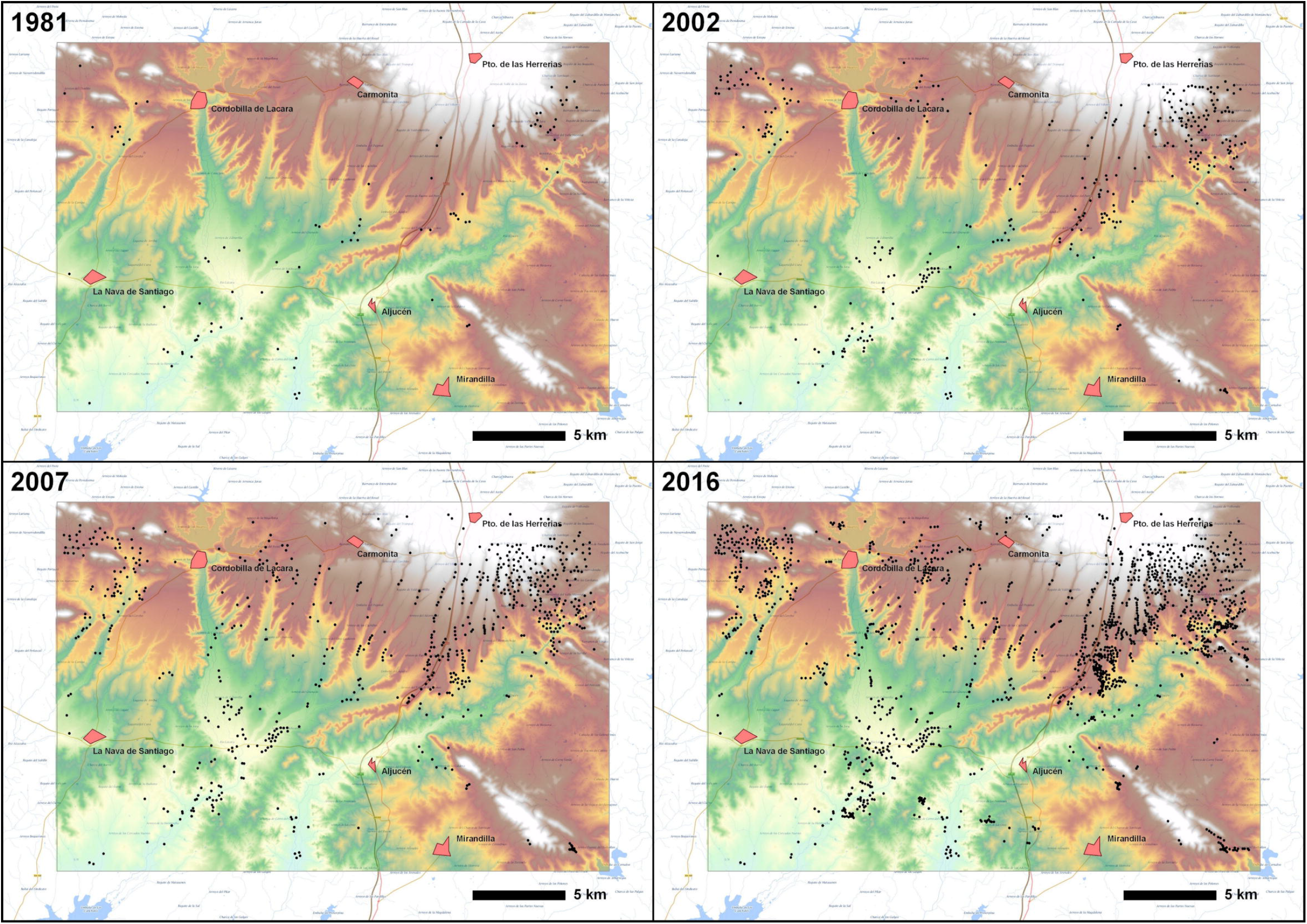
Point distributions representing the Iberian Oak Decline foci detected by computer-aided photographic interpretation of aerial images at the four different dates. Background correspond to a topographic map with elevation data represented with colours (blues and greens for lower elevations, yellows and browns for intermediates and greys and whites for the highest elevations)

**Table 1.** Aerial imagery data collections. (B&W: black and white; RGB: red-green-blue; IR infrared)

| Date | Agency | Technology | Spectral resolution | Scale | Spatial resolution |
| --- | --- | --- | --- | --- | --- |
| 1945 | US Army Map Service | Film | Panchromatic, B&W | 1:33,000 | 1.00 m |
| 1956 June | US Army Map Service | Film | Panchromatic, B&W | 1:33,000 | 1.00 m |
| 1981 March | Interministerial | Film | Panchromatic, B&W | 1:18,000 | 0.50 m |
| 1984 September | IGN | Film | Panchromatic, B&W | 1:30,000 | 1.00 m |
| 1998 December | MAPA/Oleicola | Film | Panchromatic, B&W | 1:30,000 | 1.00 m |
| 2001 August | IGN/Quinquenal | Film | Colour, RGB | 1:40,000 | 1.50 m |
| 2002 | SIGPAC | Film | Colour, RGB | 1:30,000 | 0.50 m |
| 2005 September | PNOA | Digital metric camera | Colour, RGB | 1:30,000 | 0.50 m |
| 2007 | PNOA | Digital metric camera | Colour, RGB | 1:15,000 | 0.25 m |
| 2009 | PNOA | Digital metric camera | Colour, RGB | 1:30,000 | 0.50 m |
| 2011 | PNOA | Digital metric camera | Colour, RGB+IR | 1:15,000 | 0.25 m |
| 2013 July | PNOA | Digital metric camera | Colour, RGB | 1:30,000 | 0.50 m |
| 2016 | PNOA | Digital metric camera | Colour, RGB | 1:15,000 | 0.25 m |

**Table 2.** Covariates used in the present study, including definitions and ranges. We used the variance inflation factor (VIF) with a Poisson model to quantify the severity of multicollinearity in forest stand descriptors

| Covariate | Definition | Range | VIF |
| --- | --- | --- | --- |
| Period | Time period split by dates at which images were taken | 1982 – 2002 (n= 177 foci)<br>2003 – 2007 (n= 586 foci)<br>2008 – 2016 (n= 795 foci) |  |
| FSU | Forest stand use. A binary variable describing the use of the stand (silvo-pastoral or non-wood forestry) | Forest (134 tiles, n=803, 9.3 foci/km<sup>2</sup>)<br>Open woodlands (154 tiles, n=842, 3.3 foci/km<sup>2</sup>) | 1.766 |
| TCC | Tree canopy cover (%), representing the proportion of the forest covered by the vertical projection of the tree crowns | 0 – 70 | 1.239 |
| SCC | Shrub canopy cover (%), representing the proportion of the forest covered by the vertical projection of the shrub crowns | 0 – 75 | 1.540 |
| DI | Dominance index. It indicates the degree of mixing of tree species in a stand. In balanced mixes, it takes values close to zero, as opposed to values close to 5 in pure stands | 0 – 5 | 1.085 |
| CONNECT | Stand connectivity. Categorical variable of two values: connected or isolated | Connected (265 tiles, n= 1639, 4.9 foci/km<sup>2</sup>)<br>Isolated (23 tiles, n= 9, 1.5 foci/km<sup>2</sup>) |  |

### Statistical analysis. Spatio-temporal modelling and inference

The effect of host population heterogeneities on the spread of IOD and its spatial range was analysed by means of a ‘self-exciting’ point process model continuous in space and time (Meyer et al., 2012), as implemented in the R package ‘surveillance’ (Meyer et al. 2017). In this statistical framework, the occurrence of IOD foci (represented as individual events) is modelled by the conditional intensity function (CIF) of the process. The CIF, λ*(t, s), represents the instantaneous rate or hazard for events at time t and location s, within a spatial observation window (W), given all the observations up to time t. This infection rate is modelled as a superposition of two components:

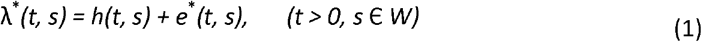

The so-called epidemic component e*(t, s) represents the spread of the disease by vectors operating at stand scale. This component makes the process ‘self-exciting’ (indicated by the asterisk) and represents the inoculum pressure due to the history of the foci cases around the point of interest. The background component h(t, s) models the rate of sporadic infections from occasional long-range transmission (possibly also from outside the study area).

In what follows, we will employ a regression method and use log-linear predictors to model both the spatio-temporal variation in the background rate (onset of new events) and the impact of the covariates on the infection pressure:

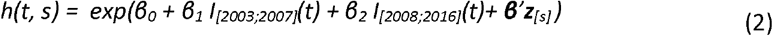

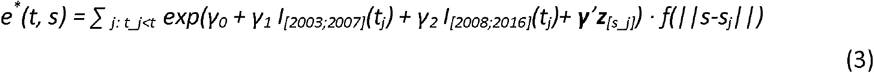

Here, the indicator functions correspond to dummy variables for the different time periods and z[s] refers to the forest stand covariates (Table 2) at location s. The epidemic component adds infection pressure from past foci j depending on the same set of covariates. The spatial interaction function (or dispersal kernel) f(x), models the decay of infection pressure with distance.

Different choices (table 3) for the spatial kernel were investigated (Meyer and Held, 2014):

**Table 3.** Mathematical equations of the spatial kernels investigated. Where ‘x’ is the distance to previous foci, ‘σ’ is a scale parameter and ‘d’ a decay parameter

| Spatial kernel | Mathematical equation |
| --- | --- |
| Constant | $f(x) = 1$ |
| Gaussian | $f(x) = \exp(-x^2/(2\sigma^2))$ |
| Student | $f(x) = (1 + x^2/\sigma^2)^{-d}$ |
| Power law | $f(x) = (1 + x/\sigma)^{-d}$ |
| Lagged power-law | $f(x) = (x/\sigma)^{-d}$ for $x \geq \sigma$ and $f(x) = 1$ otherwise |

For comparison, we also estimated a simple epidemic model with a homogeneous triggering rate, i.e., e*(t,s) = |{j: t_j_<t}| · exp(γ0), where all past foci are equally infectious regardless of their location.

Based on the typical size of fenced farms and on the home range of large mammals, we assumed the maximum distance for infection via disease vector propagation to be 2 km, corresponding to f(x) = 0 for x > 2. A temporal decay g(t-t_j_) is sometimes incorporated in such models as well but cannot be reliably estimated for the IOD spread given the limited temporal resolution of the data. Although it is known that *P. cinnamomi* propagules remain infective for up to six years (Zentmyer and Mircetich, 1966), the infectious period for an entire focus of IOD is not well characterized. Based on our own observations, we assumed this period to be longer than the time range of the study (35 years), so the model allows new foci to trigger further infections throughout the whole observation period.

Data gathered from aerial images taken at specific dates yield interval-censored infection times, where we only know the infection took place sometime during the previous observation period. To conform with a continuous spatio-temporal point process model we introduce random time shifts within the corresponding observation period and perform a corresponding sensitivity analysis, estimating each model for 60 replications of this random imputation scheme. We select the best spatial kernel based on differences in Akaike’s information criterion (AIC).

Maximum likelihood inference for the above model is implemented in the R package *‘surveillance’* (Meyer et al., 2017). We fit the model to the observed IOD pattern during the three consecutive time periods 1982-2002, 2003-2007 and 2008-2016, conditioning on the foci observed until 1981 as the so-called ‘prehistory’ of the point process. The goodness of fit of the selected model is assessed by investigating rescaled residuals in time (Ogata, 1988) and Pearson residuals on spatial pixels (Clements et al., 2011).

## Results

### Oak decline detection

Computer-aided photographic interpretation of the aerial imagery in the study area revealed a cumulative number of 1645 of cases (foci) over the whole study period. In the 1981 images, a total of 87 foci were detected. These were mainly located along the Lacara river and near the two upper corners of the boundary of the study area. In the 2002 imagery, 177 additional foci were found, most of them located in the vicinity of previously detected foci (although a new cluster of disease sites was seen to emerge in the oak populations situated between Cordobilla and Carmonita towns). In 2007 and 2016, the number of detected foci increased considerably, with a total of 586 and 795 new cases, respectively, but no new large clusters were observed.

### Phytophthora cinnamomi presence

During field visits, declining trees showing typical symptoms were observed in all the 22 stands inspected. These symptoms were like those described in earlier reports of *Phytophthora-caused* oak decline: crown transparency, branch dieback, leaf decolouration, and sudden wilting and death of entire crowns of some trees. In the sample pits, necrotic or missing fine feeder roots appeared. *Phytophthora cinnamomi* was isolated from soil rhizosphere in 29 out of 58 samples gathered during the survey (see table 4). No isolations of *P. cinnamomi* were obtained from the baits floating in downslope streams. In contrast, *Phytophthora lacustris* Brasier was recovered from all the twelve streams sampled. When subjected to a NCBI BLASTN search, the obtained ITS sequences showed a high degree of homology (94-100%) with *P. cinnamomi*. See table 4.

**Table 4.** Isolations of *Phytophthora cinnamomi* from 22 sampled sites in the study area. Indicated are the type of host species (*Quercus suber* (Qs), *Quercus ilex* (Qi) or a combination of both), the number of samples gathered at each site, and the corresponding number of *P. cinnamomi* positives. For ten sequenced isolates (regions ITS4 and ITS6), the percentage of similarity and the GenBank accession number of the best homologue accession are given.

| Site | Host | Samples | *P. cinnamomi*<br>Positives | BLAST search<br>identities (%) | GenBank accession |
| --- | --- | --- | --- | --- | --- |
| Prado de Lacara | Qs | 2 | 1 | 100 | EU079953.1 |
| Campomanes | Qs | 1 | 1 |  |  |
| Cerro Gato | Qi | 2 | 2 | 100 | EF150870.1 |
| Coto Menor de Vera | Qi | 7 | 3 |  |  |
| Coto Naharro | Qi+Qs | 5 | 1 | 100 | MN135945.1 |
| Coto Pisine | Qi+Qs | 6 | 4 | 99 | GU111591.1 |
| Cuarto de la Jara | Qi | 2 | 0 |  |  |
| Dehesa Alcuescar | Qi+Qs | 4 | 4 |  |  |
| El Machal | Qi+Qs | 3 | 1 | 100 | GU111594.1 |
| La Raposera | Qi+Qs | 4 | 1 | 100 | MK742802.1 |
| La Raposera de Abajo | Qi+Qs | 2 | 2 | 94 | MF541547.1 |
| La Raposera de Arriba | Qi | 2 | 0 |  |  |
| Las Lagunillas | Qi | 2 | 2 | 99 | EU000160.1 |
| Las Llanas | Qi+Qs | 2 | 1 |  |  |
| Las Mengachas de Arriba | Qi+Qs | 2 | 0 |  |  |
| Longueras | Qi+Qs | 1 | 1 |  |  |
| Los Machales | Qi+Qs | 2 | 1 | 99 | GU111594.1 |
| Navavaca | Qi | 2 | 2 |  |  |
| Rincon de Gavilanes | Qi+Qs | 4 | 0 |  |  |
| San Cristobal | Qi+Qs | 2 | 0 |  |  |
| Tierra Gorda | Qi+Qs | 1 | 1 |  |  |
| Valderrey | Qi | 2 | 2 | 100 | MN135945.1 |

### Spatio-temporal model output: Model components

The AIC comparison (table 5) shows that a model with background component only was outperformed by the more realistic models including an epidemic component, even with a homogeneous triggering function (Δ AIC = - 632.7). The addition of forest stand descriptors as epidemic covariates resulted in a moderately better fit (Δ AIC = - 14.9) of the two-components model.

**Table 5.**
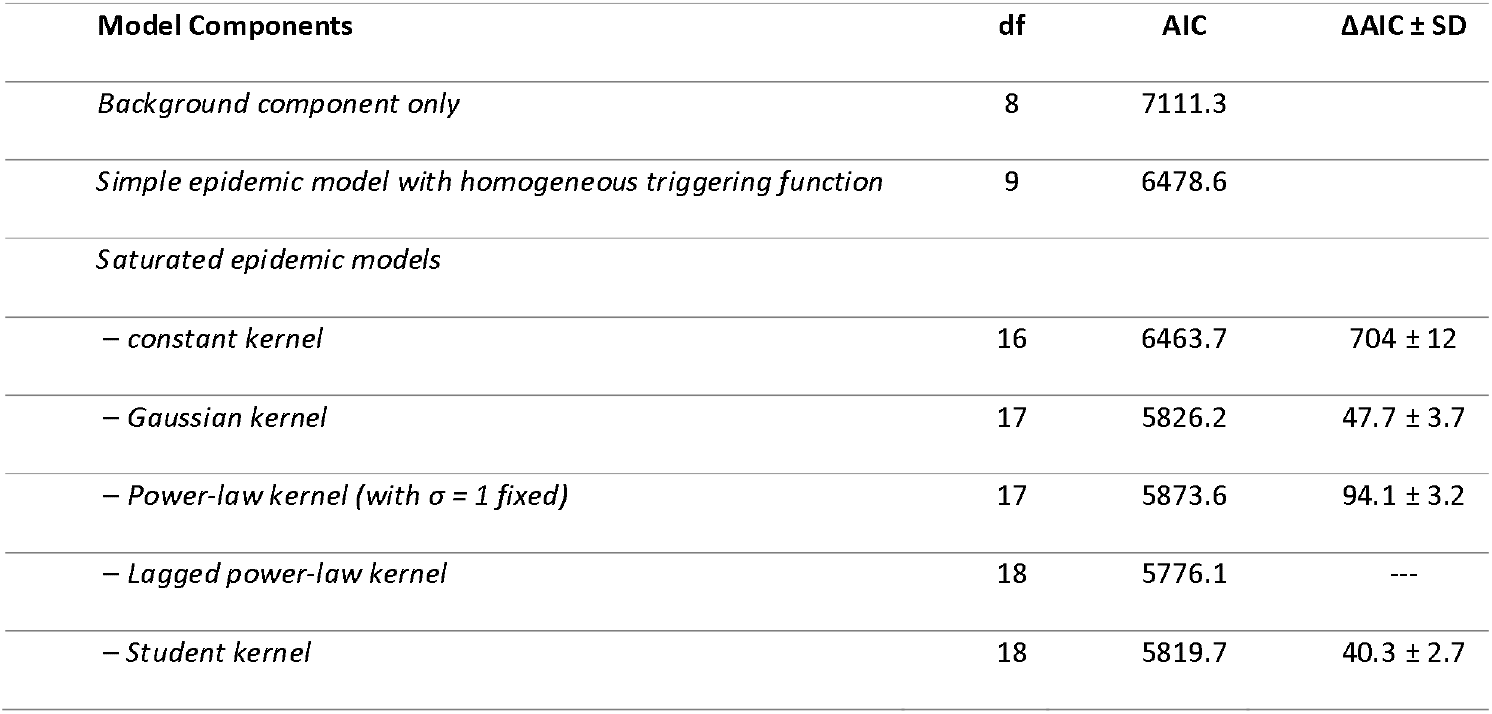
Relative model performance based on Akaike’s Information Criterion (AIC). The last column shows mean AIC differences (ΔAIC) with standard deviation (SD) of competing spatial kernels compared to the best-fitting kernel, based on 60 replications of a random imputation scheme.

| Model Components | df | AIC | ΔAIC ± SD |
| --- | --- | --- | --- |
| *Background component only* | 8 | 7111.3 |  |
| *Simple epidemic model with homogeneous triggering function* | 9 | 6478.6 |  |
| *Saturated epidemic models* |  |  |  |
| – *constant kernel* | 16 | 6463.7 | 704 ± 12 |
| – *Gaussian kernel* | 17 | 5826.2 | 47.7 ± 3.7 |
| – *Power-law kernel (with $\sigma = 1$ fixed)* | 17 | 5873.6 | 94.1 ± 3.2 |
| – *Lagged power-law kernel* | 18 | 5776.1 | --- |
| – *Student kernel* | 18 | 5819.7 | 40.3 ± 2.7 |

Introducing a spatial kernel to account for a decay of infectivity with increasing distance from the infection source yielded large improvements in AIC, most pronounced for the lagged power-law kernel. The ranking of the different kernels relative to the lagged power-law was confirmed by the sensitivity analysis, where the Student kernel ranks second (mean ΔAIC = 40.3), and the Gaussian kernel third (mean ΔAIC = 47.7). For the standard power-law kernel we had to fix σ = 1 for identifiability, which gave a fit much worse than the Gaussian kernel (ΔAIC = 94.1 compared to lagged power-law).

### Background risk factors

Table 6 shows rate ratios of covariate effects from the AIC-optimal model with a lagged power-law. The IOD background rate has changed considerably over time: compared to the baseline period 1982-2002, the background rate was estimated as 23-fold during the second period 2003-2007, and as 4-fold during the final period 2008-2016. It is likely that this saturation effect was due to the natural constraints of the ecosystem (e.g., natural barriers such as mountains, riverbeds, etc.).

**Table 6.** Rate ratios (RR) for background and epidemic effects in the AIC-optimal model with lagged power-law kernel. The table shows rate ratios (RR), i.e., exp-transformed coefficient estimates with 95% Wald confidence intervals (CI), and associated p-values. A rate ratio of 1 means there is no effect of the covariate at hand.

| Model component | Covariate | RR | 95% CI | p-value |
| --- | --- | --- | --- | --- |
| Background | *Period 2002-2007* | 23.048 | 16.16 - 32.87 | <0.0001 *** |
|  | *Period 2007-2016* | 4.359 | 2.76 - 6.89 | <0.0001 *** |
|  | *Forest Stand Use (open woodlands)* | 0.377 | 0.26 - 0.55 | <0.0001 *** |
|  | *Tree Canopy Cover (steps of 10%)* | 1.291 | 1.17 - 1.43 | <0.0001 *** |
|  | *Shrub Canopy Cover (steps of 10%)* | 1.294 | 1.13 - 1.40 | 0.0002 *** |
|  | *Dominance Index (1-5)* | 0.887 | 0.82 - 0.96 | 0.0031 ** |
|  | *Connectivity (Isolated)* | 0.352 | 0.11 - 1.18 | 0.0900 |
| Epidemic | *Period 2002-2007* | 1.582 | 1.21 - 2.07 | 0.0009 *** |
|  | *Period 2007-2016* | 2.813 | 2.22 - 3.56 | <0.0001 *** |
|  | *Forest Stand Use (open woodlands)* | 1.254 | 0.99 - 1.54 | 0.0600 . |
|  | *Tree Canopy Cover (steps of 10%)* | 1.085 | 1.01 - 1.17 | 0.0034 ** |
|  | *Shrub Canopy Cover (steps of 10%)* | 1.053 | 0.94 - 1.18 | 0.3500 |
|  | *Dominance Index (1-5)* | 1.011 | 0.95 - 1.07 | 0.7300 |
|  | *Connectivity (Isolated)* | 1.507 | 0.51 - 4.45 | 0.4600 |

Signif. codes: 0 '\*\*\*' 0.001 '\*\*' 0.01 '\*' 0.05 '.' 0.1 ' ' 1

The background IOD rate in silvo-pastoral oak stands turned out to be 62.3% (RR = 0.377; 95% CI: 0.26-0.55) lower than in forest stands. Tree and shrub cover were also associated with the background rate of IOD: a 10% increase of either tree or shrub cover (ceteris paribus) was estimated to yield a 30% increase of the background rate. The model also estimates the IOD rate to decrease with the dominance index, that is, with increasing prevalence of one oak species over the other (RR=0.887; 95% CI: 0.82--0.96). There was no evidence for an effect of connectivity.

### Epidemic risk factors

The rate of contagion was estimated to increase by 60% during the period 2003-2007 compared to 1982-2002. The infection rate during the subsequent 2008-2016 period was nearly three times the baseline value from the period 1982-2002 (see table 6). Except for tree canopy cover, no other covariates appeared to be associated with the epidemic rate. A 10% increase in oak canopy cover resulted in an 8.5% increase of the epidemic rate (RR = 1.085; 95% CI: 1.01-1.17).

### Model-based effective reproduction numbers R0

Using the AIC-optimal model, we estimated the expected number of secondary foci originating within 2 km from a primary IOD focus over 30 years. Assuming the rate from the final period 2008-2016, a focus located in a forest (TCC= 40, SCC= 10, DI= 2) is estimated to give rise to 3 secondary foci on average (95% CI: 2.4 - 3.7). Similar R0 values were obtained for foci located in silvo-pastoral dehesas with less canopy cover (TCC= 10, SCC= 0 and DI= 2), where the average number of offspring per foci is estimated to be 2.8 (95% CI: 2.0 - 3.9).

### Interaction functions

The form of a lagged power-law was confirmed by estimating a non-parametric step function with knots at 200 m, 325 m, 450 m, 575 m, 700 m, and 825 m (figure 5a). At short distances from primary foci (< 200 m), the best-fitting model is characterized by a uniform spread (figure 5b, 5c). Between 200 and 600 m (figure 5c), the lagged power-law exhibits a fast decay, which is even more pronounced than the decay of the Gaussian fit. Indeed, the estimated power-law kernel suggests that 49% of the infections triggered by an infected site are expected to occur within 250 m. For distances greater than 600 m, the power-law function exhibits a heavier tail than the Gaussian kernel, implying a higher frequency of occasional transmissions over larger distances (figure 5d).

**Figure 5.**
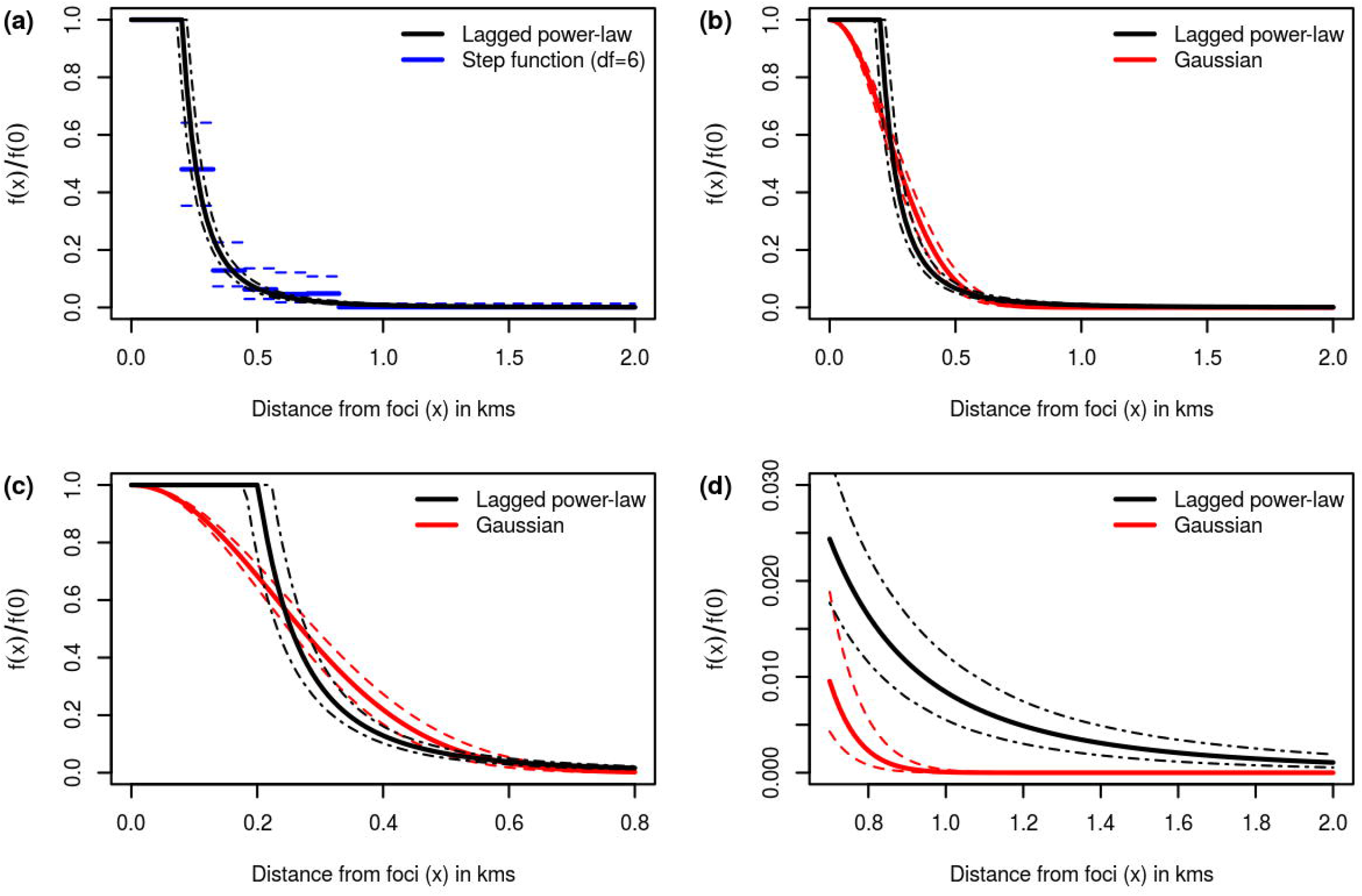
Comparison of the estimated lagged power-law kernel (f(x) = (x/σ)-d for x ≥ σ, and f(x)=1 otherwise) with the step and Gaussian kernels. a) Lagged power-law vs. step function b) Lagged power-law vs. Gaussian kernel. c) and d) Detailed plots showing the comparative behaviour of the Gaussian fit vs. the lagged power-law fit for x<800 m and x>800 m, respectively. The interaction functions have been “standardized” by division by the reference value at zero distance from the source. The dashed lines represent pointwise 95% Wald confidence intervals.

### Disease intensity and risk component weights

The spatial distribution of the disease intensity, defined as the number of foci per km^2^ accumulated over time (figure 6a), is not uniform. The map shows a very high concentration of foci in the North-East quadrant, where the intensity estimated by the model reached values almost ten times higher than the average (4.6 foci/km^2^). High intensities were also found in the North-West direction and along the south stretch of the Lacara River. We also calculated the epidemic proportion of the accumulated intensity as a function of the spatial location, i.e., which part of the intensity can be explained by local contagion. Figure 6b shows that short-range contagion was the predominant hazard (proportion > 0.8) over almost the entire study region. However, a large region of approximately 5000 hectares with very low infectivity and a low disease intensity could be observed in the South-East.

**Figure 6.**
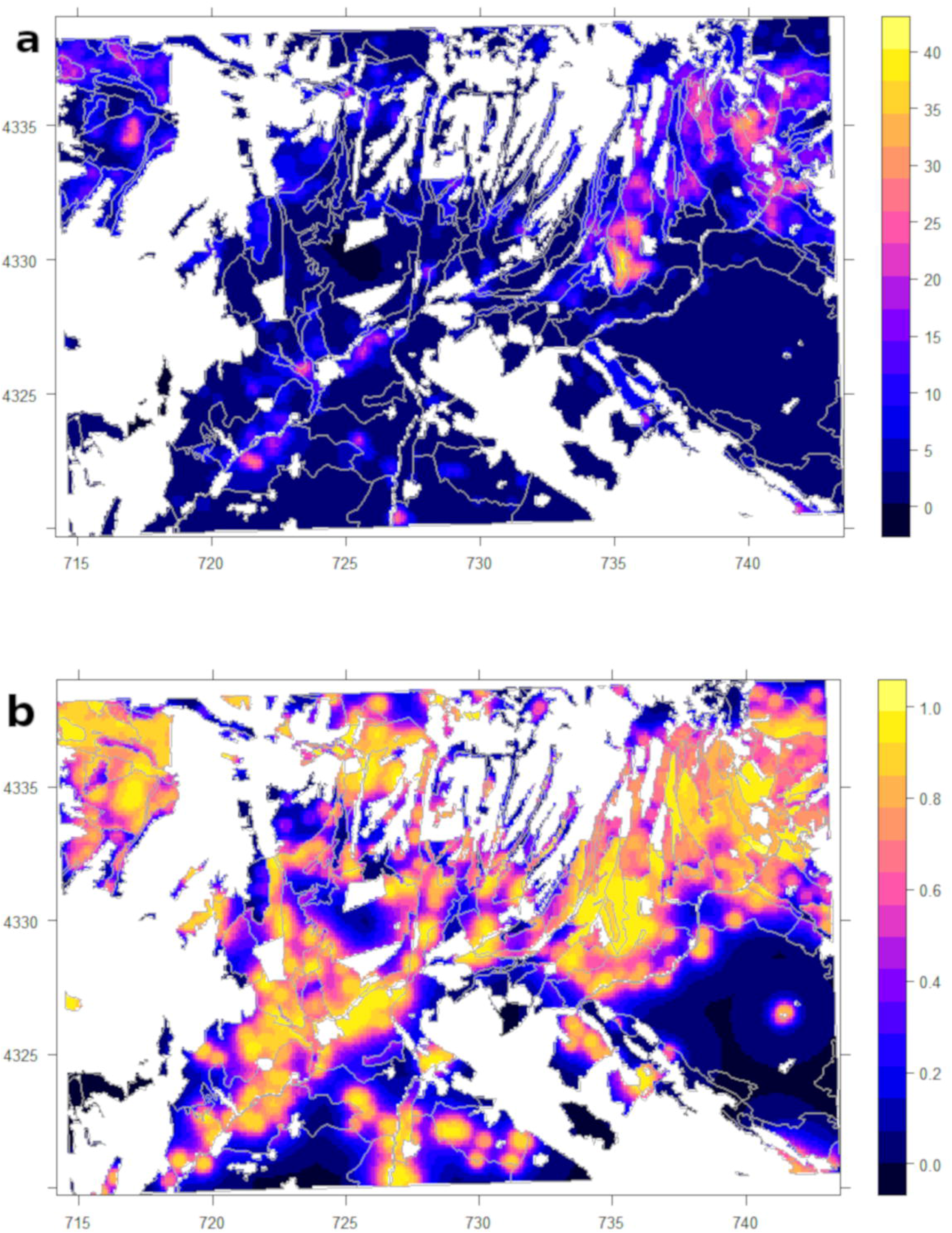
a) Model-based estimation of the disease intensity (accumulated number of foci / km^2^) of Iberian Oak Decline in the study area. b) Proportion of the epidemic intensity among the total disease intensity explained by the point process model. UTM coordinates in kms referenced to the WGS-84 system

### Goodness of fit

The graphical checks for goodness of fit do not object the suitability of the selected point process model; in particular, the estimated spatio-temporal conditional intensity function seems to provide a good description of the observed IOD spread. The transformed temporal residuals show neither evidence for lack of uniformity (fig. 7a) nor serial correlation (Fig.7b). Furthermore, integrating the estimated CIF over time, spatial Pearson residuals by 500 m x 500 m pixels (Fig. 8) do not show systematic regional model deficiencies.

**Figure 7.**
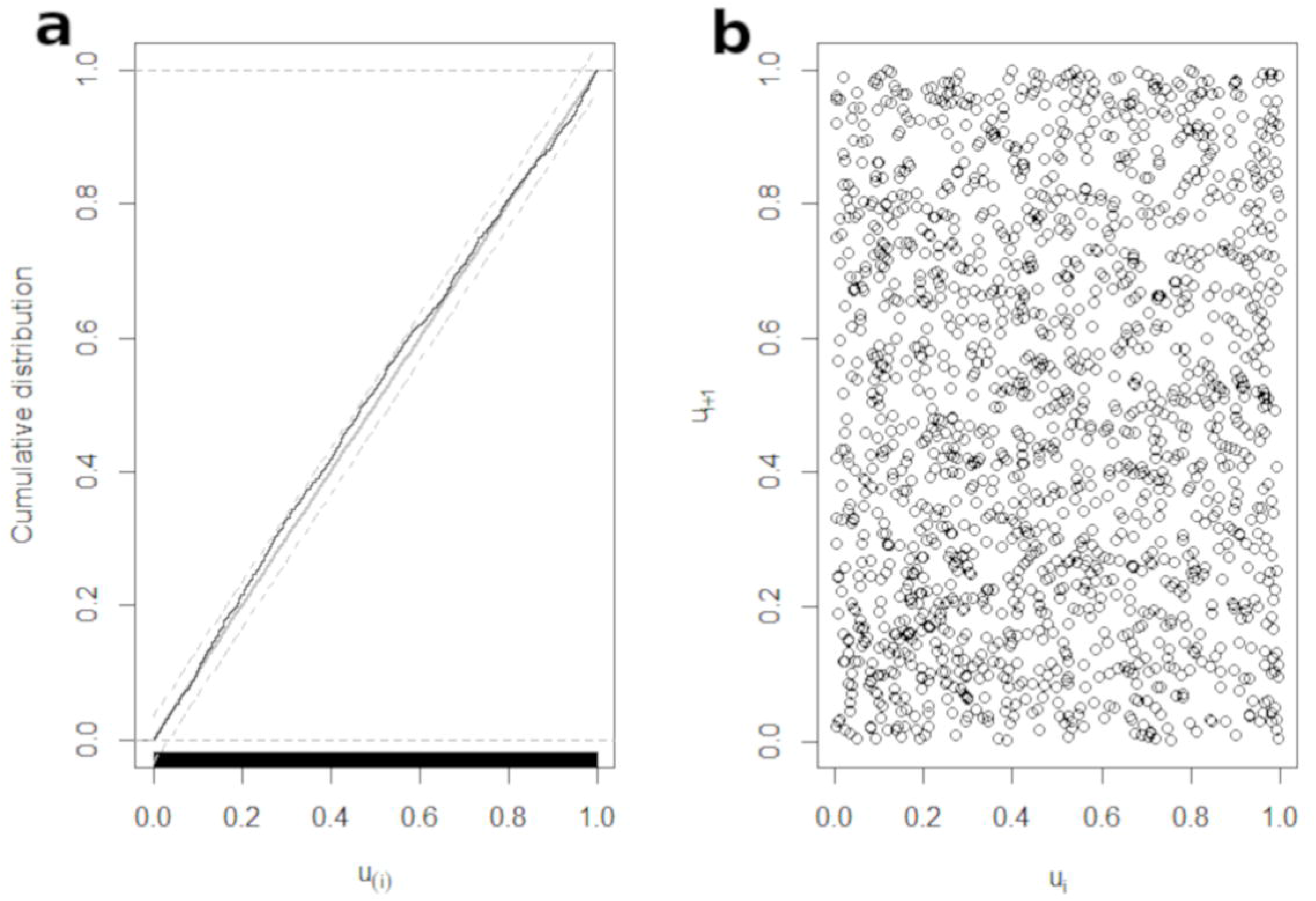
Graphical check of the goodness of fit of the temporal ground intensity estimated by the selected point process model. a) The plot on the left-hand side shows the empirical cumulative distribution function of transformed residuals (Ogata, 1988), as well as the expected uniform CDF (under the true model) with a 95% confidence band obtained by inverting the corresponding Kolmogorov-Smirnov test. No evidence for lack of uniformity of the transformed residuals is seen. b) The scatter plot on the right-hand side suggests absence of serial correlation between the transformed residuals.

**Figure 8.**
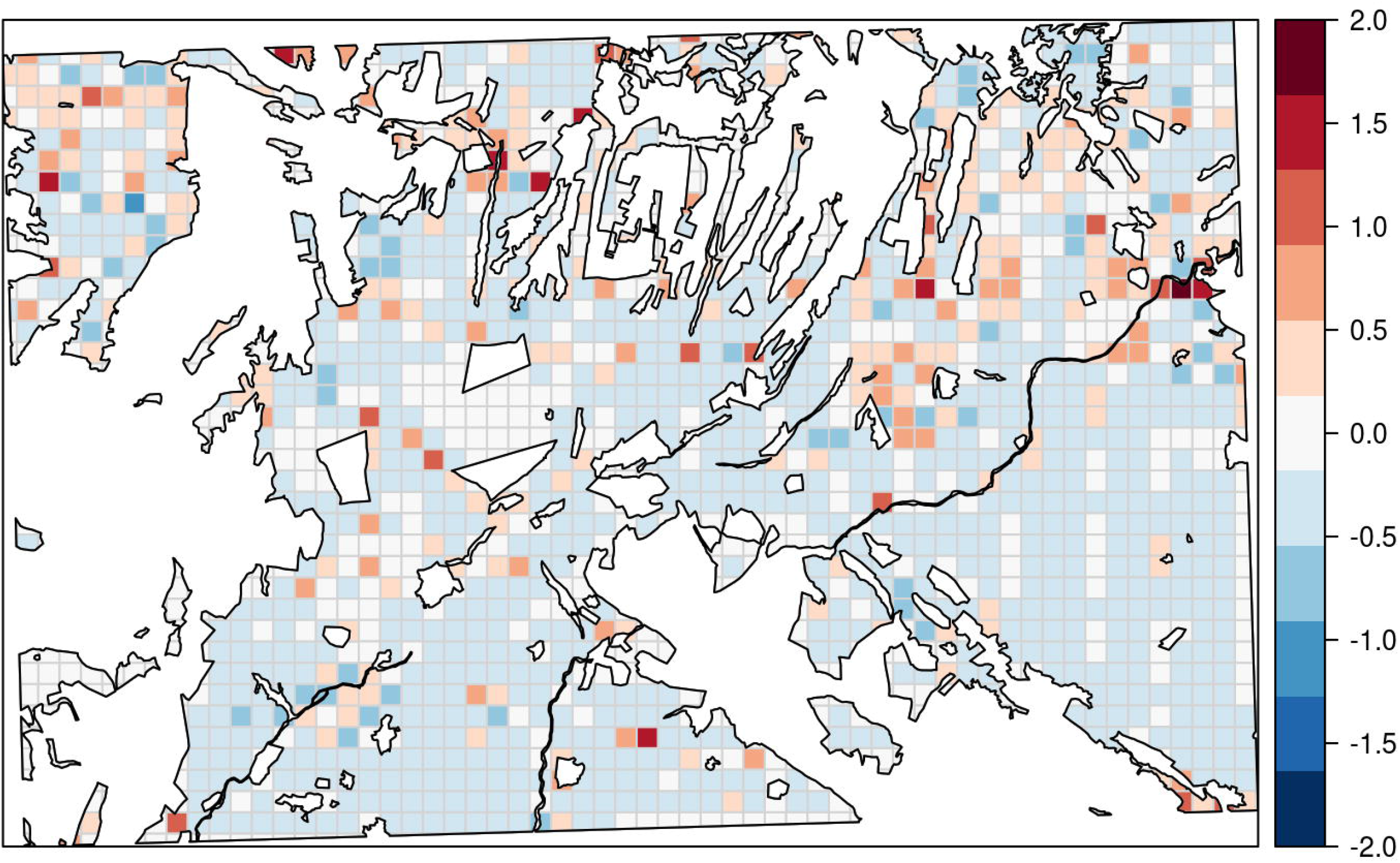
Spatial Pearson residuals, after integrating the estimated conditional intensity function over time. The grid size is 500 m x 500 m. Positive values indicate pixels where the local number of infections exceeds the corresponding model prediction.

## Discussion

We studied the spread of Iberian oak decline in forests and open woodlands of Extremadura (Spain) at landscape scale. Our purpose was to attain a better understanding of the spread patterns at landscape scale and to study the role of the host population structure in this context. To this end, we analysed the spatio-temporal dynamics of IOD by means of a ‘self-exciting’ point process model. Taking disease foci as epidemiological units, we fed the model with the locations of the foci (disease events) detected in the 33895 hectares of oak forest and woodlands at four different dates between 1981 and 2016.

### Pathogen isolation

As in other previous works (Brasier 1996; Sanchez et al. 2002; Moreira and Martins, 2005; Rodríguez Molina, et al. 2005; Serrano et al. 2012; Corcobado et al., 2013), we found a consistent relationship between the presence of *Phytophthora cinnamomi* and the onset of IOD in the study area. While we were able to obtain a high rate of isolations by rhizosphere sampling in declining and dying oaks at the surveyed sites, we failed to isolate *P. cinnamomi* by baiting downslope water streams. This failure may be due to this soil pathogen being outcompeted by other *Phytophthora* species that are more prevalent in rivers. This hypothesis is supported by the fact that we could isolate *P. lacustris* in samples from stream baits.

### Disease intensity

The photointerpretation of high-resolution aerial imagery taken at the beginning of the 1980’s reveals that, already by that time, the disease was well established (87 foci) and widely present over the entire site. The lack of high-resolution aerial photography from earlier times prevents one from estimating the precise date of disease onset; however, the exhaustive aerial imagery survey taken at three later dates provided a clear evidence for the developing forest epidemic, with 1645 accumulated foci until 2016. Admittedly, this corresponds to an epidemic of a rather low average incidence rate (0.14 foci/km^2^/year). However, it is important to point out that the disease intensity was largely inhomogeneous across the landscape, implying that large areas in which the disease was nearly absent coexisted with ravaged areas characterized by a very high cumulative disease intensity (>30 foci/km^2^).

### Time evolution

The model showed that both risk components (background and epidemic) have increased during the study period 1981-2016. We recall that the background rate during the period 2003-2007 was larger than in the initial period by a factor of 23; in addition, during the period 2008-2016, the rate of secondary foci almost doubled compared to 2003-2007. More favourable environmental conditions for inoculum establishment or reproduction could be involved in this risk acceleration. For example, it has been suggested that an increase in the number of occasional periods of unusually warm weather during the rainy season results in warm and wet soils that favour *Phytophthora* infection (Brasier, 1996; Desprez-Loustau et al., 2006). Furthermore, increased aridity is likely to have negatively affected the endurance of trees and could thus have played a significant role in the mortality process (Natalini et al., 2016), possibly exacerbating root disease caused by *P. cinnamomi* (Brasier, 1996).

Furthermore, it is known that pathogen transport mechanisms have become increasingly efficient over time. Many of the activities associated with suspected vector agents involved in *P. cinnamomi* long-range transmission were strongly intensified during the period of reference. Among these are traffic of vehicles and machinery, transport of animals, plant trade, construction of infrastructures (e.g., cattle ponds and rural roads), inspection and maintenance of farms, tourism, etc. (Moreno and Pulido, 2009). Other activities associated with transmission at shorter distances (< 2000 m), such as livestock density, big game populations and motorised human activities in farms, have also grown in number over the last 60 years (Moreno and Pulido, 2009; Lopez-Sanchez et al., 2018). In particular, the abundance of wild boar (Sus *scrofa* L.) and red deer (*Cervus elaphus* L.) in SW Spain has been increasing at an alarming rate during decades (Bosch et al., 2014; Acevedo et al., 2014), to the extent that the density of both species has become very high at some specific sites in Extremadura (Nieto-Remedios et al., 2017).

### Epidemic risk

We found that IOD has a strong epidemic character over most of the study area; indeed, a very high percentage (> 80 %) of the estimated intensity turned out to stem from the epidemic component (see fig 6b), which strongly emphasizes the prevalence of short-range (< 2000 m) dispersion. More specifically, the estimated power-law suggests that 49% of secondary foci are to be expected within less than 250 m from pre-existing inoculum sources. This characteristic range agrees well with previous estimates for other *Phytophthora* epidemics. For example, in Oregon tanoak forests infected by *Phytophthora ramorum*, Hansen et al. (2008) found that 79% of the new detections of infected trees occurred within a 300 m radius from trees that had been previously identified as diseased.

The analysis of the spatial interaction function (kernel), that is, the frequency distribution of the distance of the secondary foci from a given inoculum source (primary focus), was shown to play a key role in the mechanism of spread. Assume, for example, that the empiric kernel for vector mediated IOD transmission is well described by a Gaussian distribution; under this premise, the central limit theorem ensures that the motion of the relevant disease vectors occurs by Fickian diffusion (Skellam, 1951; Shigesada and Kawasaki, 1997). Gaussian kernels have indeed been widely used as part of fundamental models aiming to describe a large variety of propagation phenomena in many fields, including ecology and epidemiology (Shigesada and Kawasaki, 1997).

In contrast, the best fit to the actual evolution of the epidemic in order was obtained with a lagged power-law kernel, that is, a kernel combining a constant (maximal) infectivity over the first 200m with a pronounced (yet heavy-tailed) decay at larger distances. In other words, in comparison with the Gaussian model, the dispersal is enhanced at short distances, but at the same time it is also more skewed to longer distances. This strongly suggests that at least two kinds of vectors (respectively accounting for short- and long-range dispersal) are involved in IOD epidemic propagation.

This is not entirely surprising, since heavy-tailed kernels have been widely used to model stratified dispersal in which short- and long-range transmission modes are simultaneously at play (Hengeveld, 1989; Dybiec et al. 2009). Stratified dispersion often makes the disease propagation faster since, in addition to neighbourhood contagion, new colonies (in our case, pathogen sources or foci) are created at distant sites populated with healthy, susceptible individuals. As a result of this, such individuals fall within reach of the short-range infection mechanism; consequently, secondary infection foci grow, and the global amount of pathogens available for occasional long-range dispersal also increases, whereby the propagation process becomes self-sustained (Shigesada and Kawasaki, 1997).

In the above context, oak death propagation in northern Californian forests caused by *Phytophtora ramorum* has been recently shown to be compatible with epidemic models including standard power-law dispersal kernels [see supplementary material in Filipe et al. (2012)]. In our case, however, the best fit is given by a lagged power-law kernel rather than by a standard power-law. Among other effects, the initial plateau of the power-law might reflect the fact that there is a sharp transition in the propagation conditions of the pathogen beyond a certain cut-off distance. The distance x is measured from the centre of each focus; since the majority of trees inside an already existing focus are already infected, one expects a relatively low number of detectable new infections (the areal number must actually be somewhat higher, since new infections inside a focus with a high population of symptomatic individuals are very difficult to detect from the imagery because of the limitation set by the image resolution). However, as soon as x becomes larger than the typical focus radius (61 m, sd=19m), there is a sharp increase in the empirical infectivity, followed by a progressive decay at even larger distances. The net effect of the behaviour of the infectivity as a function of x over (roughly) the first 200m seems to be well captured by the plateau of the lagged power law. This may be one of the reasons why it works better than a standard power law. Another reason for the transition from spatially homogeneous spread to a power-law tail at larger x may be the presence of fences/property boundaries, which constrain the motion of some vectors in a rather effective way. Finally, approximating foci by their centroid points may distort the fitted kernel.

It is widely recognised that, in addition to very short-ranged mechanisms, such as root-to-root contact and soil water transport of propagules, (Ristaino and Gumpertz, 2000; Cardillo et al., 2018), *Phytophthora* can be dispersed via foot traffic of cattle, wild animals or humans, as well as by vehicles carrying infected soil (Ristaino and Gumpertz, 2000; Jules et al., 2002; Keith et al., 2012). At the level of the spatial scale relevant for the present work (landscape), both short- and long-range vectors are sufficiently often at play to be considered meaningful transmitters.

As already mentioned, in comparison with the Gaussian distribution, the best-fitting kernel is compatible with an enhanced frequency of infections within 200 m from primary foci. This suggests that the contribution of foot traffic is more important than contributions from other sources, such as vehicles. In the latter case, the deposition of pathogen propagules typically occurs at longer distances from the source. Interestingly, the study area is mainly composed of fenced estates with an average size of about 200 hectares. The fences are rather strong barriers for livestock and human displacements (the typical distance between fences is 800 m). However, most of these fences, while impenetrable for cattle, can be crossed by wild animals. Admittedly, other types of short-range displacements between neighbouring properties are possible, but much less frequent. Examples involving such displacements are traffic on community roads, veterinary services, cattle feed supplies, tourism activities involving human trafficking, or patrolling forest rangers.

In general, we found no relationship between stand structure and epidemic infection rates. An exception was tree canopy cover, which was found to be associated with an increased risk of shortrange infections. Host abundance has long been recognised as an important factor in the spread of forest diseases (Holdenrieder et al., 2004; Ostfeld and Keesing, 2012; Prospero and Cleary, 2017). In particular, it was also found to play an important role in the dispersion of other *Phytophthora* species in forest ecosystems. For example, in Oregon, Jules et al. (2002) showed that the risk infection by *Phytophthora lateralis* increased with stand density of the Port Orford cedar, which is the main host. Host density increases the probability of root contact and of successful establishment of the pathogen stemming from the limited quantities of propagules that can reach a new site (Jules et al., 2002).

### Background risk

The background component of the point process models the rate of sporadic cases originating from unidentified sources or long-range infection. According to the model results, open woodlands stands for silvo-pastoral use were roughly 60% less prone to background infections than forest stands without cattle. Tree and shrub canopy cover also contributed to the background rate. Our model predicted a 29% rate increment for every ten per cent rise in tree canopy cover, as well as a similar increment for shrubs. This increased background risk in thicker forests is in agreement with previous results reported by Moreira and Martins (2005), who found that more than 50% of the shrub flora in oak woodlands were asymptomatically infected with *P. Cinnamomi*. In the same way Costa et al., (2010) found higher oak mortality in oak woodlands with shrub encroachment and in shrubs and oak stands than in oak woodlands without shrubs.

Increased infectivity could be due to an enhanced capability of oaks to act as hosts for inoculum establishment, to a reservoir effect, to a more favourable environment for the action of the disease vectors at play, or to a weighted combination of all these factors. At first sight, this result could seem counterintuitive, since in silvo-pastoral open woodlands, human presence and long-range transmission vectors, such as vehicles and machinery, are more abundant than in thicker and shrub encroached forest. However, it turns out that big wild ungulates use thick forest stands as shelters and corridors; therefore, the presence of these animals is usually greater than in open woodlands (Bosch et al., 2014). As a matter of fact, wild boars and feral pigs have been reported as *Phytophthora cinnamomi* vectors by several authors (Kliejunas and Ko, 1976; Krull et al. 2013; Li et al., 2014). As such animals feed on roots and bulbs digging in soil and frequently wallow in mud, they may carry infected material both internally and externally and subsequently transport this material over considerable distances. Food and water resources are usually less abundant in forest areas than in open woodlands, and this shortage forces wild animals to search for them over longer distances. This search often implies crossing estate boundaries, which are very permeable to wild boar and deer.

We also examined the effect of species diversity on disease intensity and we found an ‘amplification effect’ (Keesing et al. 2006), that is, irrespective of the nature of the dominant species (holm oak or cork oak), the background rate in mixed oak stands was higher than in pure stands. This finding was somewhat surprising, as we were actually expecting a dilution effect in mixed stands, in line with what had been reported in about 80% of previous studies addressing the relationship between disease and species diversity (Ostfeld and Keesing, 2012; Huang et al. 2016). Such a dilution effect was ascribed to holm oaks and cork oaks being substantially different in their susceptibility to *P. cinnamomi* (Camilo Alves et al., 2013).

In ecosystems where increased background risk in mixed stands had been observed, better conditions for *Phytophthora* establishment as well as an enhanced efficiency of long-range disease vectors were invoked as possible explanations (Kessing et al., 2006). It is indeed possible that the pathogen propagation conditions at wetter and warmer sites usually preferred by cork oaks became more favourable when such sites were also populated by holm oaks, which are more susceptible hosts.

On the other hand, the season of acorn mast has a longer duration in oak mixed stands than in pure ones. This extended feeding capacity is likely to increase the attractiveness of such stands for acorn foragers able to act as disease vectors; consequently, the number of visits of such foragers to mixed stands increases, resulting in a higher infectivity. Such complex epidemiologic mechanisms are indeed known to play a role in other diseases. For example, acorn masts are known to provoke mouse outbreaks; since mice can host ticks, the result is an increase in the population of the latter, which in turn act both as propagation vectors /reservoirs of the spirochete causing Lyme disease (Ostfeld et al., 2001).

At larger scales, one would also expect the onset of other dilution effects related to habitat fragmentation, notably at sites where isolated oak forest stands are surrounded by agricultural crops (Huang et al., 2015; Prospero and Cleary 2017). However, no evidence for differences in the respective rates for connected and unconnected stands were found here. It appears that the 23 (8%) isolated stands in the area is a low number of events to supply enough statistical power to detect such differences and therefore a broad scale study with a wider presence of isolated stands is needed here.

## Conclusion

For the ecosystem at hand, we were able to establish that the majority of new IOD infections occurred in the neighbourhood of pre-existing foci. As expected, the associated epidemic risk was found to increase with growing oak stand density.

Our findings also strongly suggest that a small, yet non-negligible fraction of the infections took place via long-range dispersal, which significantly contributed to accelerate IOD spread. However, it is somewhat surprising that the best fit does not correspond to a standard power-law kernel, but rather to a lagged power-law kernel. We hypothesize that the success of the lagged power law as an effective description of the actual behaviour may stem a) from the difference in the behaviour of the inferred infectivity as a focus boundary is crossed. This appears to be well captured by the plateau of the lagged-power law. b) from the constraints introduced by fences and property boundaries in the oak lands at larger distances. This is accounted for by the long-tail of the lagged power-law.

Climate change as well as an increased activity of the disease vectors (two possibly interrelated factors) are suspected to underlie the observed increase of IOD intensity. Thicker forests turned out to be more prone to background risk than silvo-pastoral ‘dehesas’. A study of a wider scale could be a possible way to disentangle the effect of forest fragmentation from the other host effects on disease dispersal. As a soil disease, IOD is characterised by low basic reproduction numbers; in view of this fact and of the above findings, early detection and disease reduction could be enhanced by silviculture as well as by a proper targeting of possible propagation vectors.

Possible avenues for further research could exploit the fact that high-resolution aerial imagery taken at shorter time intervals will be available in the near future. Studies based on such data arrays could be carried out to shed further light on the relationship between landscape evolution and disease intensity. Finally, in order to gain further insight into the spread mechanism of IOD, one could use an improved version of the ‘self-exciting’ point process in which the epidemic component would also explicitly account for the influence of additional environmental factors, such as edaphic properties, topographic traits, etc. While the isotropic kernels used here should be viewed as an effective coarse-grained description of the pathogen propagation in a highly heterogeneous environment, it is clear that a more refined description should account for the effect of such heterogeneities. As emphasized in previous works (Cardillo et al., 2018) topographic features and soil properties might play an important role in this context. The latter are clearly influenced by weather fluctuations which one could attempt to model via the temporal part of the propagation kernel provided that a higher temporal resolution is available.

## Acknowledgements

We also appreciate the work of Pedro Antolin, and Eusebio Dorado with pathogen sampling and isolation and the help of Alba Maria Sanchez, Laura Martin and Alberto Alvarez with PCR and ITS sequencing. And last but not least, the guidance and support of Angel Felicisimo.

## References

Brasier, C.M. 1996. *Phytophthora cinnamomi* and oak decline in southern Europe. Environmental constraints including climate change. Annales des Sciences Forestieres 53, (2–3), 347–358.

Bosch, J., Mardones, F., Pérez, A., et al. 2014. A maximum entropy model for predicting wild boar distribution in Spain. Spanish Journal of Agricultural Research, 12(4), 984–999.

Camilo-Alves C.S.P., Clara M.I.E., Ribeiro N.M.C.A. 2013. Decline of Mediterranean oak trees and its association with *Phytophthora cinnamomi*: a review. European Journal of Forest Research, 132, 411–432.

Cardillo, E., Acedo, A., & Abad, E. 2018. Topographic effects on dispersal patterns of *Phytophthora cinnamomi* at a stand scale in a Spanish heathland. PloS One, 13(3), e0195060.

Clements, R.A., Schoenberg, F.P. & Schorlemmer, D. 2011. Residual analysis methods for space-time point processes with applications to earthquake forecast models in California. Annals of Applied Statistics, 5, 2549–2571.

Ciesla W.M. 2000. Remote sensing in forest health protection. FHTET 00-03, United States Department of Agriculture Forest Service. Forest Health Technology Enterprise Team, Fort Collins, CO and Remote Sensing Applications Center, Salt Lake City, UT. 266pp.

Corcobado, T., Solla, A., Madeira, M. A., et al. 2013. Combined effects of soil properties and *Phytophthora cinnamomi* infections on Quercus ilex decline. Plant and soil, 373(1-2), 403–413.

Costa, A., Pereira, H., & Madeira, M. 2010. Analysis of spatial patterns of oak decline in cork oak woodlands in Mediterranean conditions. Annals of Forest Science, 67(2), 204.

Crone, M., McComb, J.A., O’brien, P.A., et al. 2013. Annual and herbaceous perennial native Australian plant species are symptomless hosts of Phytophthora cinnamomi in the Eucalyptus marginata (jarrah) forest of Western Australia Plant Pathology, 62(5), 1057–1062.

Crone, M., McComb, J.A., O’Brien, P.A., et al. 2014. Host removal as a potential control method for *Phytophthora cinnamomi* on severely impacted black gravel sites in the jarrah forest. Forest pathology, 44(2), 154–159.

Denman, S., Kirk, S.A., Moralejo, E., & Webber, J.F. 2009. *Phytophthora ramorum* and *Phytophthora kernoviae* on naturally infected asymptomatic foliage. EPPO bulletin, 39(1), 105–111.

Desprez-Loustau M.L., Marcais B., Nageleisen L.M., et al. 2006 Interactive effects of drought and pathogens in forest trees. Annals of Forest Science, 63, 597–612.

DiLeo, M.V., Bostock, R.M., & Rizzo, D.M. 2009. *Phytophthora ramorum* does not cause physiologically significant systemic injury to California bay laurel, its primary reservoir host. Phytopathology, 99(11), 1307–1311.

Dybiec, B., Kleczkowski, A., & Gilligan, C.A. 2009. Modelling control of epidemics spreading by long-range interactions, Journal of the Royal Society Interface, 6, 941–950.

Filipe, J.A.N., Cobb, R.C., Meentemeyer, R.K., et al. 2012. Landscape Epidemiology and Control of Pathogens with Cryptic and Long-Distance Dispersal: Sudden Oak Death in Northern Californian Forests. PLoS Computational Biology, 8(1), e1002328.

Fodor, E. 2011. Ecological niche of plant pathogens. Annals of Forest Research, 54(1), 3.

García Viñas, J.I., López Leiva, C., Villares Muyo, J.M., et al. 2006. The Forest Map of Spain 1:200,000. Methodology and analysis of general results. Forest Systems, 15(S1), 24–39.

Haas, S.E., Hooten, M.B., Rizzo, D.M., et al. 2011. Forest species diversity reduces disease risk in a generalist plant pathogen invasion. Ecology letters, 14(11), 1108–1116.

Hengeveld, R. (1989). Dynamics of biological invasions. Chapman and Hall. London. 160pp.

Huang, Z.Y.X., Van Langevelde, F., Estrada-Peña, A., et al. 2016. The diversity–disease relationship: evidence for and criticisms of the dilution effect. Parasitology, 143(9), 1075–1086.

Holdenrieder, O., Pautasso, M., Weisberg, P.J., et al. 2004. Tree diseases and landscape processes: the challenge of landscape pathology. Trends in Ecology & Evolution, 19(8), 446–452.

Jules, E.S., Kauffman, M.J., Ritts, W.D., & Carroll, A.L. 2002. Spread of an invasive pathogen over a variable landscape: a nonnative root rot on Port Orford cedar. Ecology, 83(11), 3167–3181.

Jules, E.S., Carroll, A.L., Garcia A.M., Steenbock C.M., and Kauffman M.J. 2014. Host heterogeneity influences the impact of a non-native disease invasion on populations of a foundation tree species. Ecosphere 5(9):105

Jung, T., Blaschke, H., & Neumann, P. 1996. Isolation, identification and pathogenicity of *Phytophthora* species from declining oak stands. European Journal of Forest Pathology, 26(5), 253–272.

Keesing, F., Holt, R.D., & Ostfeld, R.S. 2006. Effects of species diversity on disease risk. Ecology letters, 9(4), 485–498.

Kliejunas J.T., Ko W.H. 1976. Dispersal of *Phytophthora cinnamomi* on the Island of Hawaii. Phytopathology 66: 457-460

Krull, C.R., Waipara, N., Choquenot, D., et al. 2013. Absence of evidence is not evidence of absence: feral pigs as vectors of soil-borne pathogens. Austral Ecology, 38, 534–542.

Li A.Y., Williams N., Fenwick S.G., et al. 2014. Potential for dissemination *of Phytophthora cinnamomi* by feral pigs via ingestion of infected plant material. Biological Invasions. 16(4): 765–774.

Madden, L.V. 2006. Botanical epidemiology: some key advances and its continuing role in disease management. European Journal of Plant Pathology, 115(1), 3–23.

Meentemeyer, R.K., Haas, S.E., & Václavík, T. 2012. Landscape epidemiology of emerging infectious diseases in natural and human-altered ecosystems. Annual Review of Phytopathology, 50, 379–402.

Meyer, S., Elias, J., & Höhle, M. 2012. A space–time conditional intensity model for invasive meningococcal disease occurrence. Biometrics, 68(2), 607–616.

Meyer, S., & Held, L. 2014. Power-law models for infectious disease spread. The Annals of Applied Statistics, 8(3), 1612–1639.

Meyer, S., Held, L., & Höhle, M. 2017. Spatio-Temporal Analysis of Epidemic Phenomena Using the R Package surveillance. Journal of Statistical Software, 77(11), 1–55.

Ministerio de Agricultura Pesca y Alimentación. 2006. Mapa Forestal de España (MFE50). http://www.mapama.gob.es/es/biodiversidad/servicios/banco-datos-naturaleza/informacion-disponible/mfe50.aspx. (accessed on 22, January, 2018).

Moreira, A.C., & Martins, J.M.S. 2005. Influence of site factors on the impact of *Phytophthora cinnamomi* in cork oak stands in Portugal. Forest Pathology, 35(3), 145–162.

Moreno, G., & Pulido, F.J. 2009. The functioning, management and persistence of dehesas. In Agroforestry in Europe. Springer, Dordrecht. pp. 127–160.

Natalini, F., Alejano, R., Vázquez-Piqué, J., et al. 2016. The role of climate change in the widespread mortality of holm oak in open woodlands of Southwestern Spain. Dendrochronologia, 38, 51–60.

Nieto-Remedios FJ., Garcia-Lucas AJ., Guerrero-Prieto JC., et al. 2017. El Plan General de Caza de Extremadura. 7º Congreso Forestal Español. Sociedad Española de Ciencias Forestales. Plasencia (Spain). 7CFE01–363.

Ogata Y. 1988. Statistical models for earthquake occurrences and residual analysis for point processes. Journal of the American Statistical Association 83, 9–27

Ostfeld, R.S., Schauber, E.M., Canham, et al. 2001. Effects of acorn production and mouse abundance on abundance and *Borrelia burgdorferi* infection prevalence of nymphal *Ixodes scapularis* ticks. Vector Borne and Zoonotic Diseases, 1(1), 55–63.

Ostfeld, R.S., Glass, G.E., & Keesing, F. 2005. Spatial epidemiology: an emerging (or reemerging) discipline. Trends in ecology & evolution, 20(6), 328–336.

Ostfeld, R.S., & Keesing, F. 2012. Effects of host diversity on infectious disease. Annual review of ecology, evolution, and systematics, 43, 157–182.

Plantegenest, M., Le May, C., & Fabre, F. 2007. Landscape epidemiology of plant diseases. Journal of the Royal Society Interface, 4(16), 963–972.

Prospero, S., & Cleary, M. 2017. Effects of host variability on the spread of invasive forest diseases. Forests, 8(3), 80.

Ristaino, J.B. and Gumpertz, M.C., 2000. New frontiers in the study of dispersal and spatial analysis of epidemics caused by species in the genus *Phytophthora*. Annual Review of Phytopathology. 38, 541–576.

Rodríguez-Molina M.C., Blanco-Santos A., Palo-Núñez E.J., et al. 2005. Seasonal and spatial mortality patterns of holm oak seedlings in a reforested soil infected with *Phytophthora cinnamomi*. Forest Pathology, 35(6), 411–422.

Rodríguez-Molina, M.C., Santiago Merino, R., Blanco Santos, A., et al. 2003. Detección de *Phytophthora cinnamomi* en dehesas de Extremadura afectadas por “seca” y su comportamiento in vitro. Boletín de Sanidad Vegetal. Plagas, 29(4), 627–640.

Sanchez, M.E., Caetano P., Ferraz, J., et al. 2002. *Phytophthora* disease of *Quercus ilex* in south-western Spain. Forest Pathology, 32, 5–18.

Skellam, J. G. 1951. Random dispersal in theoretical populations. Biometrika, 38, 196–218.

Zentmyer, G.A., & Mircetich, S.M. 1966. Saprophytism and persistence in soil by *Phytophthora cinnamomi*. Phytopathology, 56, 710–712.

